# Antiviral insulin signaling during West Nile virus infection results in viral mutations

**DOI:** 10.1101/2024.06.17.599370

**Authors:** Aditya B. Char, Chasity E. Trammell, Stephen Fawcett, Laura R.H. Ahlers, Dharmeshkumar Patel, Shirley Luckhart, Alan G. Goodman

## Abstract

Arthropod-borne viruses or arboviruses, including West Nile virus (WNV), dengue virus (DENV), and Zika virus (ZIKV) pose significant threats to public health. It is imperative to develop novel methods to control these mosquito-borne viral infections. We previously showed that insulin/insulin-like growth factor-1 signaling (IIS)-dependent activation of ERK and JAK-STAT signaling has significant antiviral activity. Continuous immune pressure can lead to adaptive mutations of viruses during infection. We aim to elucidate how IIS-signaling in mosquitoes selects for West Nile virus escape variants, to help formulate future transmission blocking strategies. We hypothesize that passage of WNV under activation of IIS will induce adaptive mutations or escape variants in the infecting virus. To test our hypothesis, WNV was serially passaged through *Culex quinquefasciatus* Hsu cells in the presence or absence of bovine insulin to activate IIS antiviral pressure. We sequenced WNV genes encoding for E, NS2B, NS3, and NS5 and identified variants in *E* and *NS5* arising from IIS antiviral pressure. In parallel to the genetic analyses, we also report differences in the levels of virus replication and Akt activation in human cells using virus passaged in the presence or absence of insulin. Finally, using adult *Culex quinquefasciatus*, we demonstrated the enhancement of immune response gene expression in virus-infected mosquitoes fed on insulin, compared to control. Notably, virus collected from insulin-fed mosquitoes contained a non-synonymous mutation in *NS3*. These results contribute towards achieving our long-term goal of manipulating mosquito IIS-dependent antiviral immunity to reduce WNV or other flavivirus transmission to mammalian hosts.

## INTRODUCTION

West Nile Virus (WNV) is a flavivirus belonging to the Japanese encephalitis serogroup. The first case of WNV in North America was recorded in New York in 1999 (WNV-NY99). Within 10 years the virus had spread across the continent^1^. This pathogen is transmitted through birds and the bite of infected mosquitoes, and climate change is expected to exacerbate its threat. As global temperatures rise, the latitudes at which the virus be harbored in mosquitoes, and the ranges these mosquitoes can inhabit, will expand^2^. Currently, only a small percentage of infected persons develop the deadly neuroinvasive form of this disease. The emerging threat of WNV is that millions of people will be at risk of infection in the near future. It is necessary, therefore, to increase our understanding how this pathogen interacts with its vectors and hosts to combat this threat. Using the *Drosophila melanogaster* model, a novel antiviral immune pathway in mosquitoes was elucidated that is stimulated by ingested mammalian insulin^3^. This pathway is also relevant in human cells where it interacts with endothelin signaling to recruit antiviral immunity^4^. This study presented here is aimed towards understanding how this antiviral mechanism in the mosquito vector affects the genome and virulence of WNV.

*D. melanogaster* has been previously used as a model organism to show that insulin-mediated induction of the Janus kinase/signal transducer and activator of transcription (JAK/STAT) signaling plays a vital role in host survival and immunity to WNV. We have reported that insulin-mediated JAK/STAT signaling is conserved in *Culex quinquefasciatus,* a significant vector of WNV^3^. JAK/STAT signaling is mediated by the domeless receptor cascade in insects^5^. Briefly, the binding of endogenous insulin or insulin-like peptides (ilps) to insulin-like receptor (InR) triggers the phosphorylation of Akt and Extracellular Signal-Regulated Kinase (ERK). Phosphorylated ERK upregulates the expression of the protein upd2/3, which in turn activates the domeless receptor. The domeless receptor signals through hopscotch and leads to activation of the transcription factor Stat92E that induces antiviral effector genes, such as *vir-1* and *Vago2*^3,6^. In parallel, the phosphorylation of Akt further causes the phosphorylation of the forkhead transcription factor FoxO. This prevents FoxO from being localized to the nucleus. The nuclear localization of FoxO induces the expression of *Dicer 2* (*Dcr2*), *Argonaute 1* (*AGO1*), and *AGO2* components of the RNAi pathway^7,8^. Thus, stimulation of InR activates JAK/STAT signaling while suppressing RNAi.

Mosquito InR is able to recognize a family of peptides called insulin-like peptides (ilps), in addition to mammalian insulin^9,10^. The ingestion of insulin has been found to reduce the titer of WNV in infected flies and mosquitoes^3^. Mosquitoes lack an adaptive immune response, and they primarily control viral infection by the RNA interference (RNAi) pathway^3,11^. The ingestion of insulin by mosquitoes stimulates antiviral genes through JAK/STAT signaling while suppressing the RNAi response^12^. Understanding how JAK/STAT affects virus mutation could be important in developing methods to control this pathogen. This type of mitigation strategy has been suggested for developing combinational antiviral therapies for HIV. Mathematical models were used to simulate the relationship between HIV intra-host genotypes and antiviral drugs. These models would predict the optimal time to change the antiviral drug to avoid the development of resistance and virus escape^13^. A similar approach might be used to predict patterns of mutation in flaviviruses.

The effects of anti-viral immune pathways on the evolution of flaviviruses is well documented. Viral antigens recognized by host immune responses can change or the viruses develop ways of hindering these immune responses. For example, the NS5 protein of the NY99 strain of WNV (WNV-NY99) is able disrupt JAK/STAT signaling. This activity is linked to a single peptide change compared to the Kunjin NS5 protein, which shows weak JAK/STAT antagonism^14^. WNV-NY99 can also delay IRF-3 signaling which is an important transcription factor for innate immune responses^15^. The diverse range of hosts that the *Flaviviridae* family infect places unique pressure on the evolution of these viruses. The genomes of viruses that infect only vertebrate or invertebrate hosts can be distinguished by their unique codon usage and dinucleotide patterns^16^. In mosquitoes, the RNAi immune response drives WNV diversification as rare genotypes that can escape sequence-specific short interfering RNAs^17^. Given that the ingestion of insulin suppresses this pathway, WNV will be subjected to a novel selective pressure from IIS-mediated immunity.

This study aims to catalogue the impact of the insulin-mediated antiviral immune response on the genome of the NY99 strain of WNV. The immune response stimulated by the insulin receptor in insects such as *D. melanogaster* has been shown to improve survival during infection with the Kunjin strain of WNV^3^. In the work presented here, we demonstrate that IIS-mediated selection is equally significant against the pathogenic WNV-NY99 strain. Key components of IIS, including the insulin receptor and ilp7, improve survival during infection in *D. melanogaster*. Additionally, insulin stimulation was found to reduce WNV-NY99 load in insect cells *in vitro* as well as in adult flies. To investigate the impact of continuous IIS-mediated selection pressure on WNV, we serially passaged WNV-NY99 through *Culex* Hsu cells and performed sequenced WNV genes after each passage to catalog genetic changes. Under IIS selection pressure, we identified unique viral genotypes whereby WNV appeared to lose its capacity to manipulate Akt signaling in mammalian cells. *In vivo* infection in *Culex quinquefasciatus* under IIS pressure resulted in a mutation in the NS3 viral protein, with molecular dynamics simulations predicting impaired NS3 protein function. Our findings reveal that WNV under IIS-mediated selection pressure develops distinctive genotypic and phenotypic characteristics, highlighting the importance of this antiviral immune pathway during WNV infection in the mosquito vector.

## RESULTS

### Insulin reduces WNV-NY99 infection in *D. melanogaster* and insect cell lines

In insects, InR, ilp, and their downstream signaling pathways have multiple evolutionarily conserved functions including larval development, regulation of feeding and immune response regulation^18,19^. To demonstrate the importance of IIS in *D. melanogaster* during WNV-NY99 infection we knocked-down *InR* by RNAi and used mutant flies deficient in *ilp7*. Previous studies have found that these genes were similarly important for survival against the attenuated WNV-Kun strain^3^. Unlike other fruit fly ilps, ilp7 is conserved in mosquitoes^19^. We infected control and *InR* knockdown or *ilp*^7^ mutant *D. melanogaster* with 2,000 PFU of WNV-NY99 by intrathoracic injection and monitored survival for 30 days. These results show that the knockdown of InR significantly increased the susceptibility of flies to WNV-NY99 infection (**Fig. 1A**). Similarly, mutant flies lacking *ilp*^7^ were more likely to succumb to WNV-NY99 infection (**Fig. 1B**).

**Figure 1.**
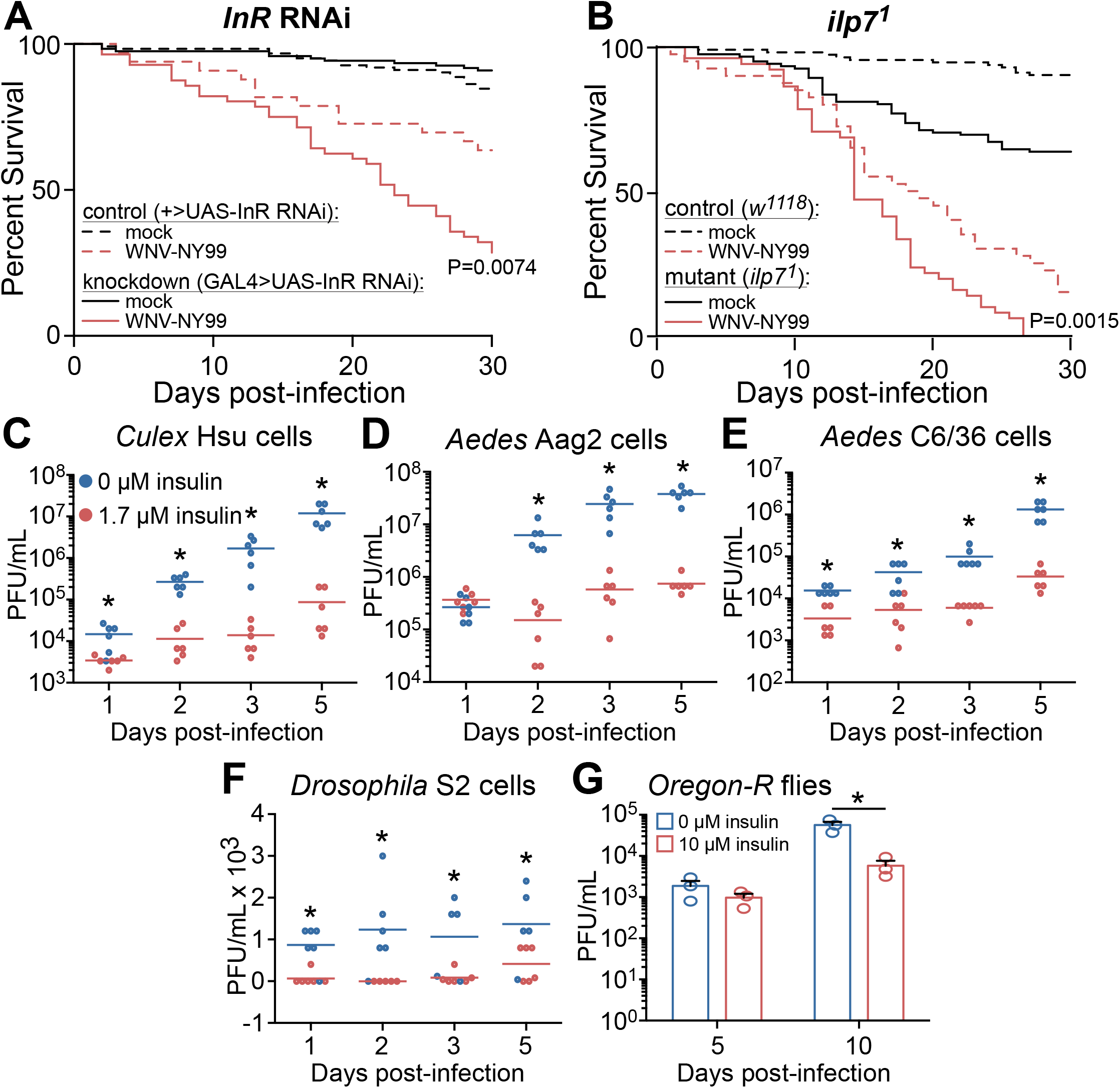
Insulin signaling protects against mortality to WNV-NY99 infection and reduces virus replication. (A) *InR* knockdown and (B) *ilp7*^1^ mutant *D. melanogaster* are more susceptible to WNV-NY99 (2,000 PFU/fly) compared to genetic controls. Significance compared to infected control *D. melanogaster* is indicated (Log-rank test). (C-F) WNV-NY99 (MOI=0.01 PFU/cell) titer is reduced in (C) *Cx. quinquefasciatus* Hsu, (D) *Ae. aegypti* Aag2, (E) *Ae. albopictus* C6/36, and (F) *D. melanogaster* S2, cells primed with 1.7 μM bovine insulin for 24 h prior to infection. (G) WNV-NY99 (dose 2,000 PFU/fly) replication is reduced in adult *D. melanogaster* fed 10 μM bovine insulin (Unpaired t-test; *p<0.05).

To show the importance of insulin signaling during WNV-NY99 infection in insect cells, we treated multiple mosquito cell lines and *D. melanogaster* S2 cells with 1.7 μM bovine insulin prior to WNV infection. Similar to previous findings using WNV-Kun^3^, we found that insulin treatment reduced WNV-NY99 viral load throughout infection (**Fig. 1C-G**). We showed that the antiviral activity of IIS-mediated immunity is functional in mosquito cell lines from *Culex* (**Fig. 1C**) and *Aedes* species (**Fig. 1D-E**), as well as *Drosophila* S2 cells (**Fig. 1F**). We next tested if ingested insulin reduced WNV-NY99 infection in adult *D. melanogaster*. Fly food was supplemented with 10 μM bovine insulin or vehicle control. By 10 dpi, flies in the insulin-fed group had lower levels of WNV compared to control flies (**Fig. 1G**). Taken together these data show that insulin signaling reduces levels of the pathogenic strain of WNV in adult flies and insect cells, supporting our previous results using the Kunjin strain of WNV^3^.

### Mutations arise during serial passage of WNV-NY99 in insulin-treated *Cx. quinquefasciatus* Hsu cells

The negative effects of innate immune systems on virus growth can induce the development of viral evasion strategies^20^. For instance, the receptor binding domain (RBD) of SARS-CoV-2 was reported to develop escape mutations when the virus was cultured with anti-RBD antibodies^21^. Similarly, the envelope protein of HIV is known to develop mutations in conserved sites targeted by therapeutic broadly neutralizing antibodies^22^. Individual immune pathways have exert unique selective pressures in comparison to these pathways working in concert^23^. We hypothesized that serial passage of WNV during insulin stimulation would select for virus sequence variants, since insulin stimulates JAK/STAT immunity while suppressing the RNAi response^3^. We used serial passage of WNV-NY99 in Hsu cells under sustained insulin-mediated selection, with cells infected with virus from the previous passage for 10 passages. Cells were primed with media containing 1.7 μM bovine insulin or vehicle control 24 hours prior to infection with virus. Virus load was analyzed at each passage. We observed that insulin treatment significantly reduced the virus load in Hsu cells in 7 of the 10 passages (**Fig. 2**). Both groups showed a drop in titer at passage 5 followed by recovery at later passages. Specifically, after the drop in titer at passage 5, both groups had significantly higher titer at passage 8. Following passage 8, however, control WNV remained at high titer while the insulin-treated group yielded virus titers at passages 9 and 10 that were similar to those of passage 5. To test if sustained insulin-mediated selection was associated with changes in the WNV-NY99 genome, we performed Sanger sequencing of WNV genes *E, NS2B, NS3,* and *NS5* in virus collected from infected cells from passages 5 and 9 (**Table 1**). The proteins coded by these genes play important functions during the establishment of the disease. The E protein is an important structural protein that is a major target of the host immune response^24^. The proper conformations of this protein in many WNV variants, including NY99, require an N-linked glycosylation at residues 154-156. A study examining this glycosylation site reported that mice infected with WNV-NY99 mutants lacking this motif were less likely to develop the neuroinvasive disease^25^. The NS3 protein is a multi-functional molecule and plays many important roles in WNV infections. It has helicase activity, nucleotide triphosphatase activity and in conjunction with NS2B, it acts as a serine protease^26^. This protease cleaves the other non-structural proteins from the viral polyprotein during infection. The NS2B/NS3 protease of Dengue virus, another flavivirus, is known to disrupt IFN type I production^27^. The NS5 protein of WNV-NY99 is known to be a potent antagonist of type I interferon activity. Research comparing the NS5 proteins of WNV-NY99 and WNV-Kun showed that the NY99 protein was 30-fold more effective at suppressing IFN-mediated JAK/STAT signaling^14^.

**Figure 2.**
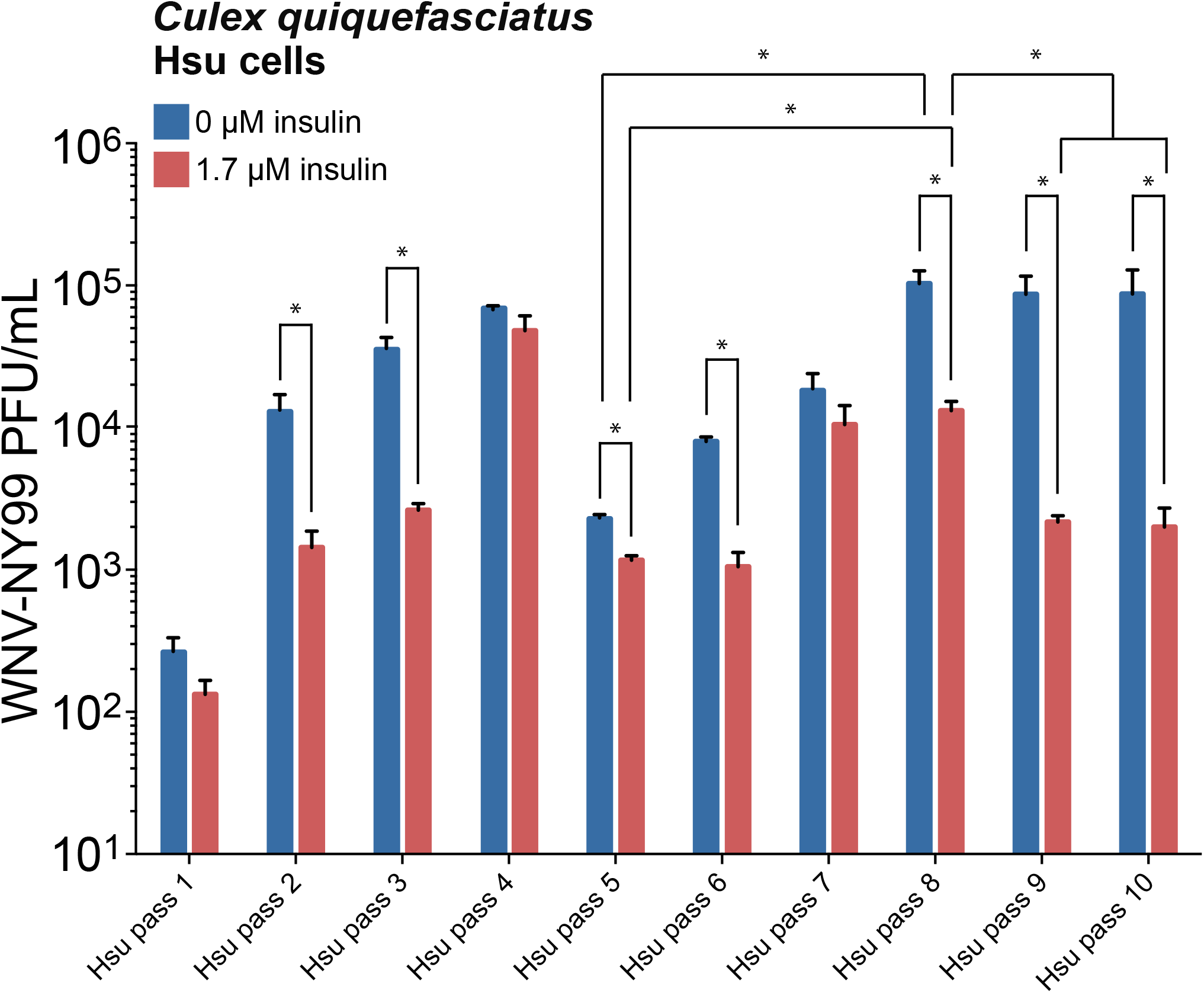
Replication of WNV-NY99 serially passaged in Hsu cells. Hsu cells were treated with 1.7 μM insulin or vehicle control, then infected with WNV-NY99 (MOI=0.001 PFU/cell). Media from infected cells was collected 5 dpi. Titer was determined by plaque assays. Data are the average titer of three biological replicates (Unpaired t-test; *p < 0.05).

**Table 1.** Codon Usage Analysis of WNV-NY99 *Envelope* (*Env*) and *NS5* Genes.

| Treatment | Passage | Gene | SNP Position on consensus | Codon Change (WT/SNP) | Silent mutation? | Amino acid | Codon usage fraction in *Culex* (SNP/WT) | Codon usage fraction in *H. sapiens* (SNP/WT) |
| --- | --- | --- | --- | --- | --- | --- | --- | --- |
| 1.7 μM insulin | Pass 5 | *Env* | C1603A | cuC/cuA | Yes | Leucine | 0.61 | 0.37 |
| 1.7 μM insulin | Pass 5 | *Env* | C549U | guC/guU | Yes | Valine | 1.22 | 0.76 |
| 1.7 μM insulin | Pass 5 | *NS5* | G1861A | cuG/cuA | Yes | Leucine | 0.16 | 0.18 |
| 1.7 μM insulin | Pass 9 | *Env* | C549U | guC/guU | Yes | Valine | 1.22 | 0.76 |

In passage 5 and 9 virus, we did not detect any mutations in the *NS2B* and *NS3* genes compared to the original stock genome. However, we did detect mutations in the *envelope* and *NS5* genes, with a single synonymous C to T transition at position 549 of the *envelope* gene that persisted from passage 5 to 9. We calculated the codon usage in *Cx. quinquefasciatus* and humans to represent the significance of the codon bias for each mutant. Synonymous codons are not used equally within organisms and this bias can affect translation rate, causing ribosome pausing while the less abundant tRNAs are recruited. This could also affect the timing and pattern of protein folding^28^. Codon usage fractions less than one represent mutant codons that are less abundant in the organism while values greater than one represent mutant codons that are more abundant^28^. Codon usage fractions were calculated using values from the Kasuza codon usage database (www.kazusa.or.jp/codon)^29^. For example, we found that the codon frequency for *envelope* C1603A and *NS5* G1861A mutations was ∼2-5-times lower than wild-type (columns 8-9), suggesting decreased tRNA codon availability for the transcript variant that could affect translation of the WNV polypeptide. Furthermore, we found that the codon frequency for the persistent *envelope* C549U mutation was 22% higher than wild-type in *Cx. quinquefasciatus*, suggesting that increased tRNA codon availability for the transcript variant could counteract insulin-mediated selection pressure (**Table 1**). Together, the results suggest that serial passage of WNV-NY99 in *Cx. quinquefasciatus* cells under insulin pressure induces mutations in the WNV genome that could affect virus replication under treatment.

We next examined human cellular responses to control WNV or insulin-selected WNV. Given that flaviviruses stimulate Akt phosphorylation to promote cell survival and abrogate pro-apoptotic signaling^30^, we infected human liver HepG2 cells with passaged WNV and used western blotting to quantify Akt activation (phosphorylation) in infected cells (**Fig. 3**). This model is relevant based on our earlier work with HepG2 cells showing that endothelin signaling and associated Akt activation are induced by insulin stimulation to control WNV replication^4^. HepG2 cells infected with control virus from passages 2, 3, 7 and 8 showed strong Akt phosphorylation at 24 hours post-infection (hpi) which was absent at 72 hpi (**Fig. 3**). We also compared Akt phosphorylation between HepG2 cells infected with control or insulin-selected WNV. In passages 2 and 3 there was little difference in Akt phosphorylation between the treatment groups at 24 and 72 hpi. However, at passages 7 and 8, while HepG2 cells infected with control WNV displayed Akt phosphorylation at 24 hpi, cells infected with insulin-selected WNV displayed lower Akt phosphorylation at 24 hpi (**Fig. 3**).

**Figure 3.**
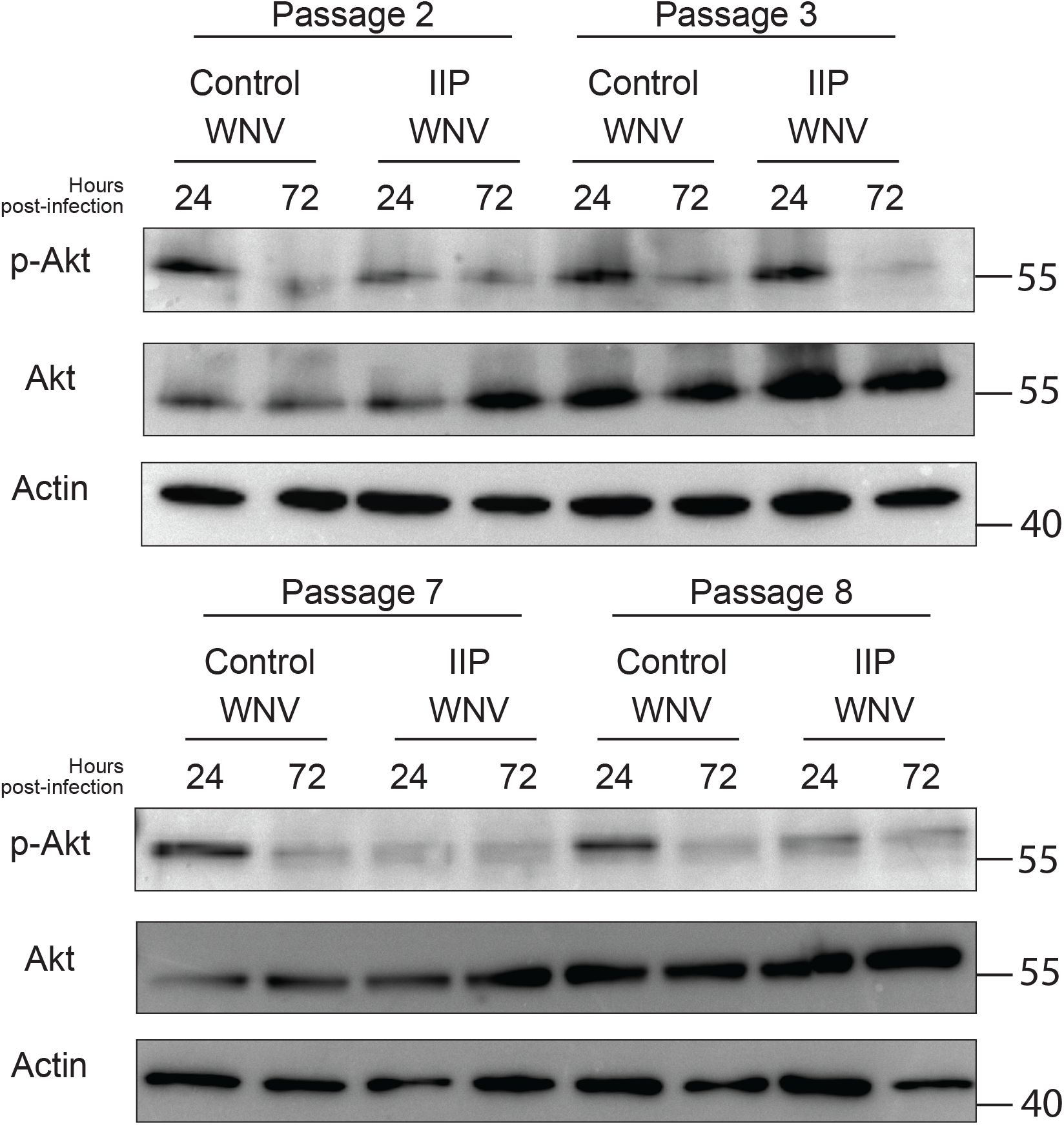
Serial passage under IIS-mediated immunity reduces Akt signaling. HepG2 cells were infected with WNV-NY99 from Hsu passage treatment groups. The supernatant from the wells of each treatment group are pooled before infecting the HepG2 cells. Cells are lysed at 24 and 72 hpi. Control WNV is collected from Hsu cells in 0 μM insulin media. IIP WNV is collected from Hsu cells in 1.7 μM media.

### Insulin activates mosquito immunity during WNV infection

Our data from Hsu cells showed that serial passage of WNV-NY99 under insulin-mediated selection was associated with mutations in the WNV *envelope* and *NS5* genes and altered host responses. Accordingly, we sought to examine host responses to infection under insulin selection in *Cx. quinquefasciatus,* the primary vector of WNV in North America^31^. We infected female mosquitoes with the Kunjin strain of WNV in the presence or absence of 170 pM bovine insulin in the bloodmeal^3^. At 10 dpi, total RNA was collected from groups of five mosquitoes in each treatment group for RNA sequencing. Using these data, we performed Gene Set Enrichment Analysis (GSEA) to analyze gene expression patterns. Gene ontology (GO) categories related to host defense and immunity^32^ were significantly more enriched in WNV-Kun infected mosquitoes provisioned with 170 pM insulin relative to controls. For example, insulin fed mosquitoes showed significant enrichment of genes in the serine-type peptidase GO category (**Fig. 4A**). Genes within this category have been linked to antiviral responses in mosquitoes against dengue virus (DENV). Multiple flaviviruses show antagonistic activity against the expression of a specific serine protease^33^. Further, the oxidoreductase category includes genes that reduce Zika virus load in mosquitoes. This GO category is highly enriched in insulin-fed mosquitoes during WNV-Kun infection^34^. We also analyzed the RNAseq data using a gene set derived from *Cx. quinquefasciatus* infected with WNV-NY99^35^ to generate a heatmap representing the fold change in gene expression compared to uninfected controls. We observed that a majority (60%) of genes were upregulated in infected mosquitoes that were bloodfed 170 pM of insulin compared to vehicle control (**Fig. 4B-C**). We next used the RNAseq results from insulin- or control infected mosquitoes to map the data to the consensus sequence for WNV-Kun (GenBank Accession KX394396.1). We identified a non-synonymous mutation in the *NS3* gene (K84I) of WNV-Kun following a single virus passage in insulin-treated mosquitoes (**Fig. 5**). The mutation was in the N-terminal protease domain close to ADP binding sites in the C-terminal helicase domain. Molecular dynamics simulations predicted that the K84I mutation affects ADP binding free energy (ΔΔG_fold_) and the free energy of the NS3-ADP complex (ΔΔG_bind_). We observed a small change in binding affinity (+0.90 kcal/mol) and observed a significant change in stability of the NS3-ADP complex (+11.10 kcal/mol). The K84 residue forms an H-bond with residue E43. A prior study examining the effects of amino acid changes on the stability of protein complexes suggested that ΔΔG values of ±2 kcal/mole and beyond represent a significant change. This value was determined by comparing free energy change predictions by simulations against experimental values^36^. The K84I mutation lost this H-bond interaction which might affect the stability of the structure. This mutation was not detected in virus from infected control mosquitoes.

**Figure 4.**
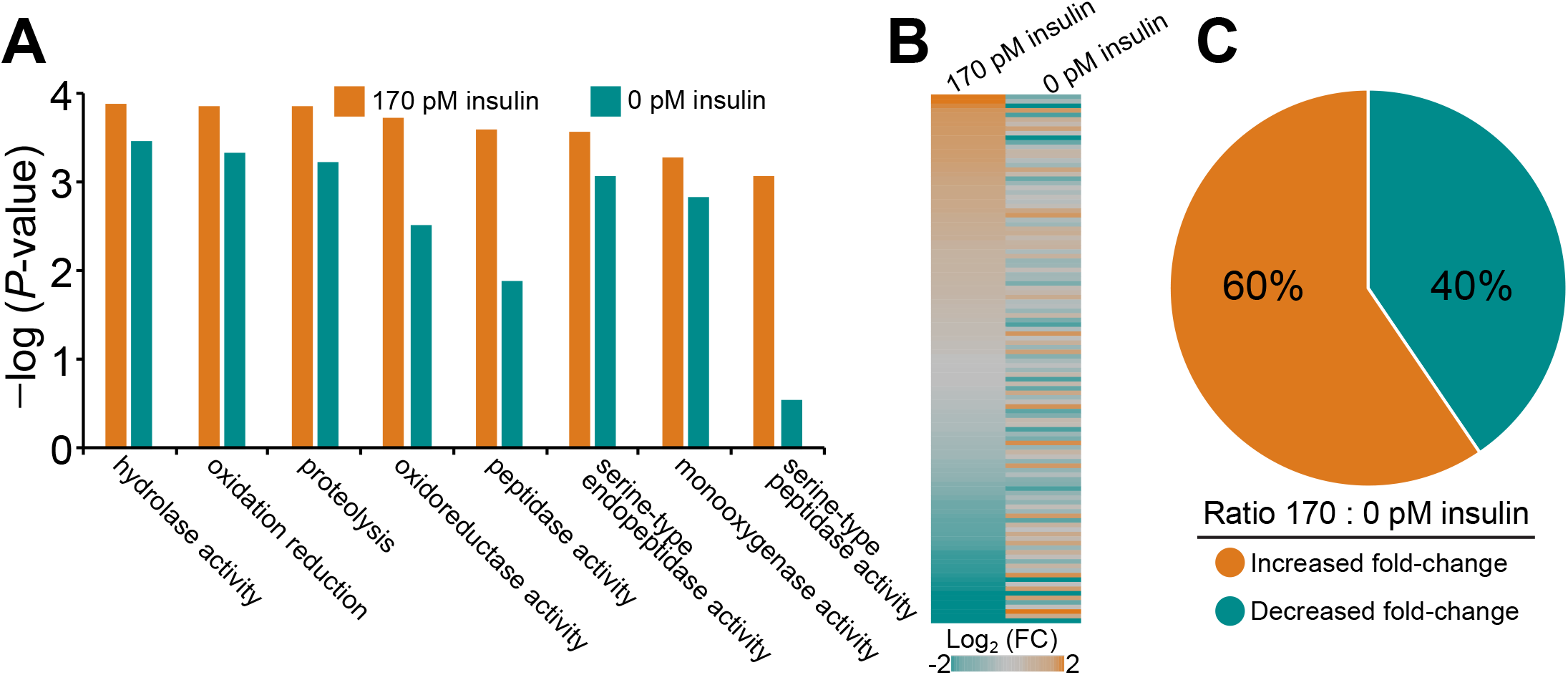
Ingested insulin induces defense responses in *Cx. quinquefasciatus*. (A) Gene set enrichment analysis (GSEA) using RNA-seq datasets from WNV-Kun-infected mock- and insulin-treated mosquitoes shows increased enrichment of defense response GO categories^32^. (B) Heatmap using geneset described previously as up-regulated during WNV-NY99 infection^35^. (C) Ratio of gene expression in (B) showing insulin up-regulates a majority of WNV-induced genes.

**Figure 5.**
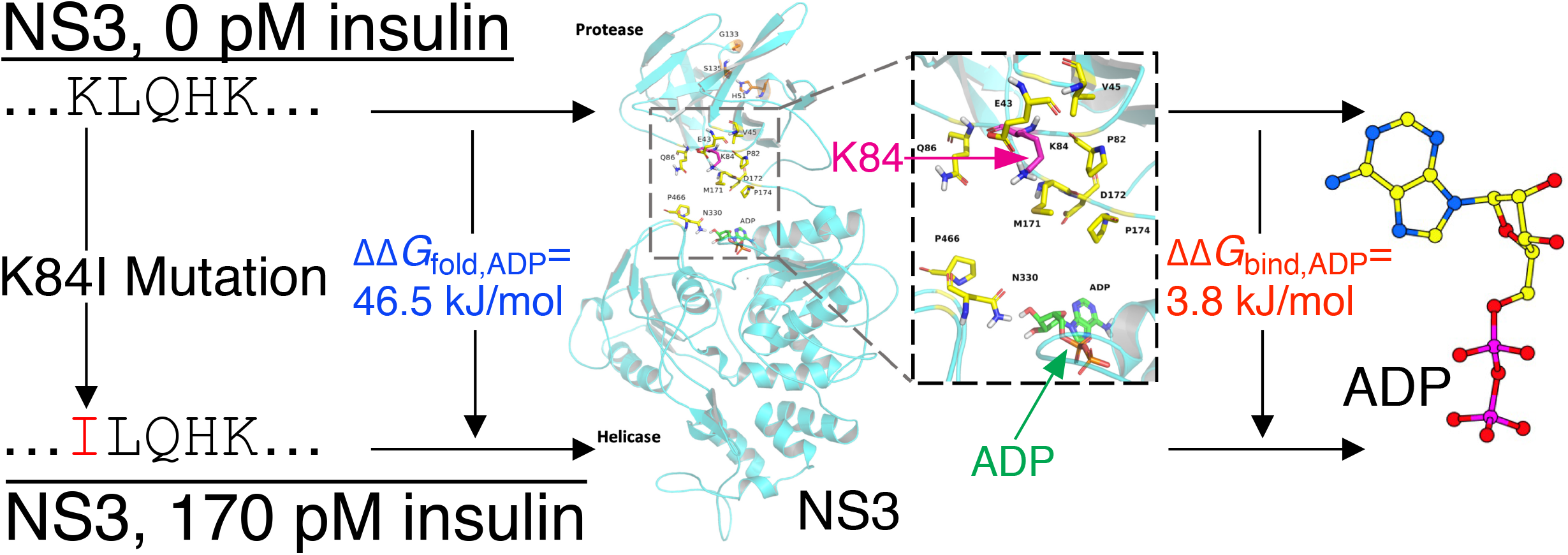
Molecular modeling of the NS3 K84I mutation. Modeling of a non-synonymous mutation in WNV-Kun following one passage in *Cx. quinquefasciatus* fed 170 pM insulin indicates that the mutant NS3 K84I protein is less stable in the presence of ADP (ΔΔG_fold,ADP_>0) and may reduce ADP binding (ΔΔG_bind,ADP_>0).

In summary, we demonstrated that IIS-mediated selection is associated with mutations in the WNV-NY99 genome both *in vitro* and *in vivo*. Additionally, insulin enhances the expression of immune genes that are predicted to contribute to host defenses against WNV. These promising findings encourage further investigation of the insulin-dependent antiviral pathways in the mosquito vector and how these shape the evolution of the WNV genome.

## DISCUSSION

Previous studies showed that insulin signaling activates important antiviral immune pathways in both *D. melanogaster* and mosquitoes^3,4,37^. Here we demonstrated that previous using the attenuated Kunjin strain of WNV hold true for the pathogenic WNV-NY99 strain as well. Notably, knockdown or ablation of components of the IIS-mediated immune pathway was detrimental to *D. melanogaster* survival during WNV-NY99 infection. Additionally, insulin priming prior to infection was found to reduce virus load in insect cell lines as well as in adult fruit flies.

The synonymous WNV mutations recorded in this study were seen in the genes coding for the envelope and NS5 proteins. The engineering of synonymous mutations in the same genes of other flaviviruses like Zika virus (ZIKV) and DENV has been associated with attenuation. For example, codon optimization was used to alter the sequence of the *envelope* gene of DENV and the *NS3* and *NS5* genes of ZIKV. These silent mutants were shown to have reduced fitness and virulence in a mosquito cell line and mice^38^. We postulate that the occurrence of silent changes in the genes of WNV over sustained insulin-mediated selection could lead to changes in virulence. We have reported multiple synonymous mutations confined to WNV under insulin-mediated selection pressure. This included a synonymous mutation in the *envelope* gene that persisted in passages 5 and 9 and a synonymous mutation in sequence of the *NS5* gene encoding the C-terminal region that functions in RNA-dependent RNA polymerase activity required for viral replication^39^.

Our data from liver HepG2 cells infected with serially passaged WNV revealed a divergence in the host response using WNV from early and late passages. Specifically, insulin-selected WNV from later passages 7-8 did not elicit Akt phosphorylation to levels observed in cells infected with control WNV from the same passages. Since late passage control WNV showed similar Akt phosphorylation levels to early passage WNV, we do not attribute this change to the serial passaging procedure itself. Accordingly, we propose that WNV manipulation of Akt signaling was lost due to sustained insulin-mediated selection on the virus genome. This would be detrimental to the virus as apoptosis of the host cell must be prevented at early stages to avoid interruption of the virus replication cycle^40^. Conversely, this loss could be caused by the physiological effects of insulin priming on Hsu cells. The application of insulin itself stimulates the phosphorylation of Akt^3^. Over the course of passaging, high levels of Akt phosphorylation induced by insulin could have supported, and even selected for, the growth of WNV variants that were unable to activate the PI3K/Akt pathway.

In this study we showed that insulin-mediated selection pressure*, in vitro and in vivo,* affects the genome of WNV-NY99. Additionally, we showed that IIS activation in the presence of WNV-NY99 infection enhances known defense responses of the mosquito vector. These data suggest that targeting IIS in the mosquito is a viable route for reducing WNV incidence and transmission in the field. There is a need for alternate approaches to WNV mitigation due to the drawbacks of currently employed strategies. For example, strategies that use *Wolbachia* to reduce viral infection in mosquitoes^41–43^ have been shown to enhance WNV infection^44^.

To support our *in vitro* studies with Hsu cells, we examined the effects of insulin provisioning on WNV in *Cx. quinquefasciatus*. A single passage *in vivo* in insulin-fed mosquitoes was associated with the appearance of a non-synonymous mutation in the WNV *NS3* gene. Our molecular dynamics simulation predicted the thermodynamic stability of the mutant NS3 K84I protein in terms of the free energy difference (ΔΔG) in the presence of ADP, compared to the wild type. The positive value of ΔΔG indicates that the mutant K84I protein has impaired ADP binding affinity while the mutant protein-ADP complex is less stable^45^. ADP plays an important role in the function of the NS3 helicase given that the translocation of the protein on RNA is ATP-dependent. The replacement of ATP with ADP, as a result of hydrolysis, would lead to the favorability of particular protein confirmation states^46,47^. A reduction in the stability of the NS3-ADP complex, therefore, could result in the mutant K84I protein being functionally sub-optimal.

Our understanding of other flaviviruses and other pathogens could benefit from the findings of this study. Despite the reduction in virus titer seen in insulin-fed mosquitoes we do not recommend the feeding of insulin itself to mosquitoes given that this has been shown to enhance mosquito infection with malaria parasites^48^. A more effective strategy could leverage the link between the insulin receptor and immune pathways to simultaneously activate RNAi and JAK/STAT signaling. The immune response produced by this combination was found to reduce the load of ZIKV in live mosquitoes to a greater degree than either pathway alone^49^.

Multiple components of the immune system are compromised in diabetic patients, leading to worse prognoses during viral infections^50^. A recent meta-analysis of studies on WNV and DENV reported that diabetes was a significant risk factor for developing severe disease^51^. Hyperglycemia has been suggested as the mechanism behind the increased risk, however the precise cause remains unclear^52,53^. The findings of this study are another step towards understanding the interplay between insulin-mediated signaling and diseases caused by WNV and other flaviviruses.

## MATERIALS AND METHODS

### Cell culture and virus

Baby hamster kidney-21 (BHK-21) (ATCC, CCL-10) and HepG2 (ATCC, HB-8065) cells were cultured in DMEM supplemented with 10% fetal bovine serum (FBS) and maintained at 37°C in 5% CO_2_. *Culex quinquefasciatus* Hsu cells were cultured in L-15 media supplemented with 10% FBS and 10% triphosphate buffer. These cells were maintained at 28°C in 5% CO_2_. West Nile virus-Kunjin strain MRM16 (WNV-Kun) was gifted by R. Tesh and propagated in Vero cells. West Nile virus strain 385-99 (WNV-NY99) was obtained from BEI Resources, NIAID, NIH (NR-158) and propagated in Vero cells. All experiments with a specific virus type utilized the same stock.

### Fly infections

Knockdown of *InR* was achieved by crossing *y^1^v^1^;P{TRiP.JF01482}attP2* (InR dsRNA cassette, Bloomington *Drosophila* Stock Center (BDSC) 31037) with *y^1^w*;P{Act5C-GAL4}25FO1/CyO,y^+^* (Actin driver, BDSC 4144). Sibling offspring not expressing the actin driver were used as controls. *Ilp7^1^* mutant flies (*w^1118^;TI{w^+mW.hs^=TI}Ilp7^1^*, BDSC 30887) were compared to *w^1118^* controls (BDSC 5905). Oregon-R-C flies (BDSC 5) were also used. Flies are negative for *Wolbachia* infection. Flies were kept on standard cornmeal food (Genesee Scientific #66– 112) at a temperature of 25°C with 65% relative humidity and a 12-hour light/dark cycle. The adult flies used for experiments were female and aged between 2 to 7 days post-eclosion. Bovine insulin (Sigma 10516) was incorporated into the food to achieve a final concentration of 10 μM. Flies were intrathoracically injected with 2,000 PFU of WNV-NY99 or PBS vehicle control, as previously described^3^.

### Serial passage

Hsu cells were seeded onto a 6-well plate at a density of 10^6^ cells per well and incubated overnight. At 24 hours post-seeding, the cell media in each well was aspirated and replaced with either 0 μM or 1.7 μM insulin media, depending on the treatment group. After 24 hours of incubation in the chosen media, the cells were infected with WNV at MOI 0.001 PFU/cell. The plates were incubated with virus for 2 hours before aspirating the infection media and replaced with 2% FBS L-15 media. Infected Hsu plates were incubated for 5 days before collection of cell media. Cells adhered to the plate were lysed in 300 μL of Trizol and stored at - 80°C to preserve RNA.

### HepG2 infection

HepG2 cells were seeded onto a 24-well plate at a density of 7.5 x 10^5^ cells per well and incubated for 48 hours. After the incubation period, the cells were infected with WNV at MOI 0.001 PFU/cell. The plates were incubated with virus for 1 hour before aspirating the infection media and replaced with 2% FBS DMEM media.

### Immunoblotting

Protein lysate from infected HEPG2 cells was collected by applying 100 μL of RIPA buffer^3^ to each well. The concentration of protein lysate was assessed by BCA assay. 5 ug worth of protein was diluted in 2x Laemmli and heated to 95°C for 5 minutes. The prepared samples were analyzed by SDS-PAGE using 10% acrylamide gel followed by transfer to a PVDF membrane. Blots used to target Akt (1:2,000) (Cell Signaling 4691) were blocked in 5% non-fat dairy milk (NFDM) in Tris-buffered saline and 0.1% Tween (TBST) before application of primary antibodies. Blots used to target P-Akt (1:1,000; Cell Signaling 4060) were blocked in 5% FBS in Tris-buffered saline and 0.1% Tween (TBST) before application of primary antibodies. Blots were incubated with primary antibodies overnight at 4°C followed by 2-hour incubation with HRP-conjugated secondary anti-rabbit IgG-HRP conjugate antibodies (1:10,000; Promega W401B) at room temperature. For actin blotting (1:10,000; Sigma A2066), PVDF membranes were stripped with gentle rocking for 30 minutes. The procedure for actin blotting was similar to Akt blots.

### Workflow for Sanger sequencing

RNA was isolated (ZymoResearch R2050) from the infected Hsu cells at various passages and 1 μg of RNA was used to prepare cDNA (BioRad 170-8891). The cDNA was used as the template for various PCR reactions targeting the 4 genes of interest - E, NS2B, NS3 and NS5 (Table 2).

**Table 2.**
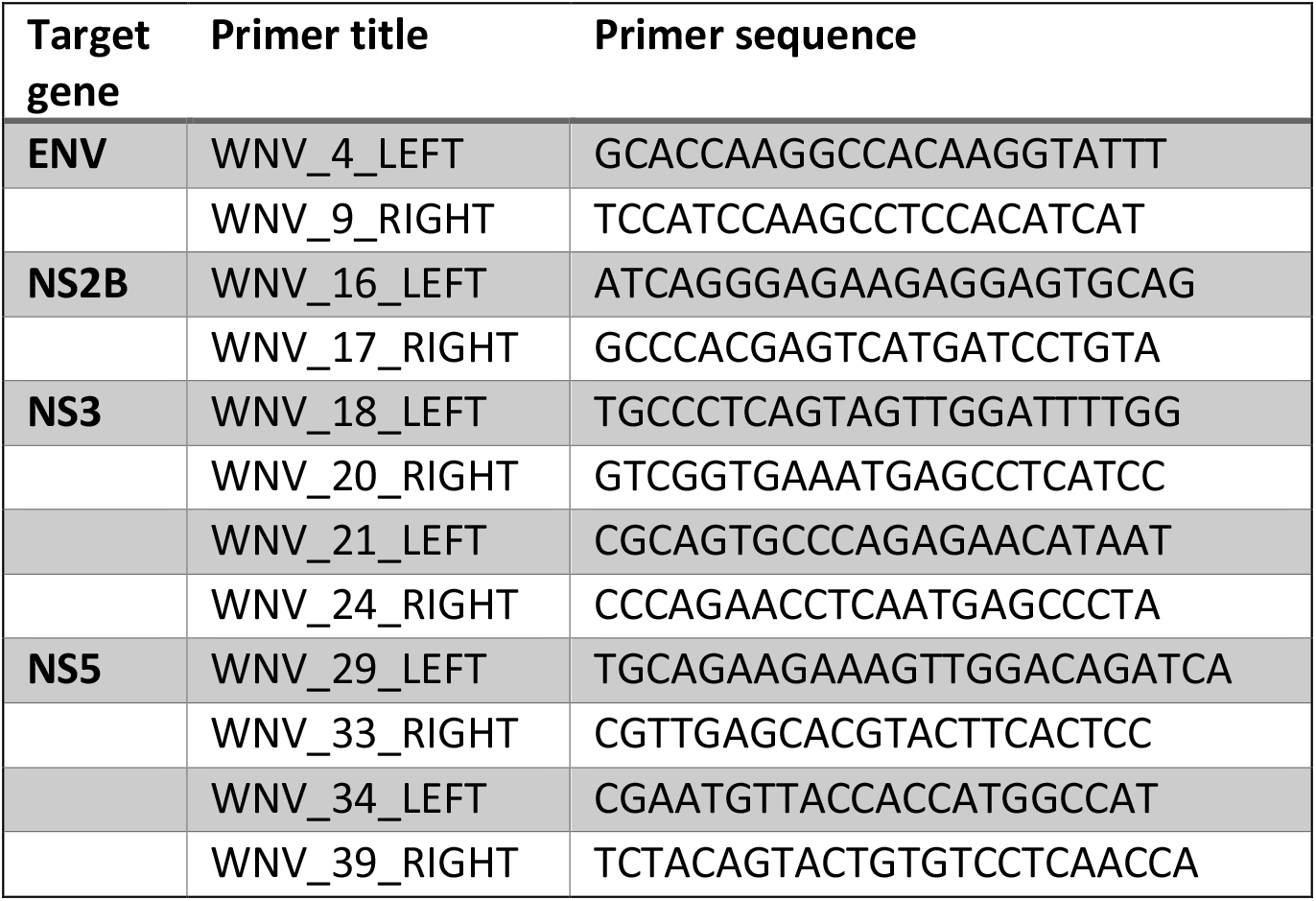
Primer list.

The genes coding for NS3 and NS5 were amplified in two parts by overlapping primer pairs. The primers targeting these genes were selected from a phylogenetic study of WNV-NY99 in Arizona which used an array of primers to sequence the whole virus genome^54^. The products from these PCR reactions were purified (NEB T1030S). The purified products were sent to Eurofins for Sanger sequencing. Forward and reverse sequencing reactions were prepared for the genes of each biological replicate. The forward and reverse sequence files were used to assemble complete gene sequences using the ‘Assemble contigs’ function in SnapGene (v7.1.1). These complete sequences were aligned to the genome of the virus used for the first passage and analyzed for mutations.

### Mosquito infection

WNV-Kun was mixed with defibrinated chicken blood that had been washed with RPMI and supplemented with 20% FBS and HEPES buffer or 170 pM insulin, described previously, resulting in a final concentration of 1×10^7^ PFU WNV-Kun/mL^48,55^. Female *Cx. quinquefasciatus* were bred for three generations and then fed using an artificial mosquito feeder (Chemglass Life Sciences) for 2 hours. Females that became engorged were separated under CO2, kept in 1-gallon cartons with constant access to 10% sucrose, and collected at 1-, 5-, and 10-days post-infection for analysis. Females that did not feed were excluded from further analysis.

### RNA extraction and RNAseq

Mosquitoes were lysed with Trizol Reagent (ThermoFisher 15596), RNA was isolated by column purification (ZymoResearch R2050) and DNA was removed (ThermoFisher 18068). Following RNA extraction, library preparation and RNAseq was performed as previously described^4^. RNA sequencing was performed at the Spokane Genomics CORE at Washington State University-Spokane in Spokane, WA, USA.

### Gene Set Enrichment Analysis

Gene Set Enrichment Analysis (GSEA) was performed using RNAseq datasets and their associated p values as previously described^56^. The GSEA used transcript levels for each experimental condition normalized to 0 µM insulin from the RNAseq analysis. GO categories were selected based on GSEA P value (P < 0.05).

### Molecular Modeling of WNV NS3

The NS3 of WNV is made up of two domains: a helicase and a protease. The crystal structure of WNV-Kun helicase is available (PDB id-2QEQ) without bound form of the ADP nucleotide. To generate the full NS3 protein, a homology model of WNV-Kun helicase-protease was generated using the structure of NS3 protease-helicase from dengue virus (PDB ID-2WHX) using the Prime-structure prediction wizard of Schrödinger (2020-4). The conformation of nucleotide ADP was transferred to the homology model from NS3 protease-helicase from dengue virus and loops of the homology model were refined using Prime Module of Schrödinger (2020-4).

Desmond, a molecular dynamics (MD) engine of Schrödinger (2020-4), was used to execute the MD calculations for predicting the stability of WNV-Kun helicase-protease-ADP complex. Two separate MD simulations were performed: 1) WNV-Kun helicase-protease-ADP complex, 2) Mutant K84I WNV-Kun helicase-protease-ADP complex. The complexes were embedded in the TIP3P water model of orthorhombic box and were neutralized with the counter ions and physiological salt concentration kept at 0.15 M of NaCl. The system was specified on periodic boundary conditions, the particle mesh Ewald (PME) method for electrostatics, a 10 Å cutoff for Lennard–Jones interactions and SHAKE algorithm for limiting movement of all covalent bonds involving hydrogen atoms. OPLS3e forcefield was used to prepare the systems.

The energy was minimized using hybrid method of steepest descent (10 steps) and the LBFGS (limited-memory Broyden–Fletcher–Goldfarb–Shanno) algorithm with a maximum of 2000 steps with solute restrained, followed by similar energy minimization for 2000 steps without solute restraints. Then 12 ps simulation was carried out in NVT (Constant volume and temperature) ensemble for restraining nonhydrogen solute atoms at 10 K temperature was repeated and then followed by 12 ps simulation in the NPT (constant pressure and temperature) ensemble of temperature 10 K for restraining nonhydrogen solute atoms. About 24 ps simulation in the NPT ensemble were restrained with solute nonhydrogen atoms at temperature 300 K and 24 ps simulation in the NPT ensemble at temperature 300 K with no restraints. Berendsen thermostats and barostats were used to control the temperatures and pressures during the initial simulations. The relaxed system was subjected to simulation time of 50 ns with a time step of 2 fs in the NPT ensemble using a Nose–Hoover thermostat at 300 K and Martyna–Tobias– Klein barostats at 1.01 bar pressure. Every trajectory was recorded with a time interval of 100 ps.

MM-GBSA was calculated for MD trajectories of both complexes 1) WNV-Kun helicase-protease-ADP complex, 2) Mutant K84I WNV-Kun helicase-protease-ADP complex using the Prime module of Schrödinger (2020-4) to calculate binding affinity (ΔG_bind_) of ADP. The difference between the former and latter provided the binding affinity change (ΔΔG_bind_) of ADP due to the K84I mutation. Residue scanning of Schrödinger (2020-4) was performed on snapshots of MD trajectory of WNV-Kun helicase-protease-ADP complex, to check the stability (ΔΔG_fold_)of the structure due to mutation K84I.

### Quantification and statistical analyses

Results shown are representative of at least three independent experiments. Data points in dot plots represent a biological replicate of an individual well of cells (Fig. 1C-F) or a pool of five flies (Fig. 1G). Statistical analyses were completed using GraphPad Prism. Two-tailed unpaired t-tests assuming unequal variance were utilized to compare normally distributed pairwise quantitative data. All error bars represent standard error of the mean. Survival curves (Fig. 1A-B) represent three replicate experiments per condition pooled together and analyzed by the log-rank (Mantel-Cox) test using GraphPad Prism to determine P values between infected genotypes.

## DATA AVAILABILITY

Raw and processed RNAseq data have been deposited in NCBI Gene Expression Omnibus (GEO) accession # GSE269077.

## ACKNOWLEDGEMENTS

This study was funded by the College of Veterinary Medicine Stanley L. Adler Research Fund, Washington State University.

## Notes

https://www.ncbi.nlm.nih.gov/geo/query/acc.cgi?acc=GSE269077

### Competing Interest Statement

The authors have declared no competing interest.

